# Quantifying state-dependent control properties of brain dynamics from perturbation responses

**DOI:** 10.1101/2025.02.18.638784

**Authors:** Yumi Shikauchi, Mitsuaki Takemi, Leo Tomasevic, Jun Kitazono, Hartwig R Siebner, Masafumi Oizumi

## Abstract

The brain can be conceptualized as a control system facilitating transitions between states, such as from rest to motor activity. Applying network control theory to measurements of brain signals enables characterization of brain dynamics through control properties. However, most prior studies that have applied network control theory have evaluated brain dynamics under unperturbed conditions, neglecting the critical role of external perturbations in accurate system identification. In this study, we combine a perturbation input paradigm with a network control theory framework and propose a novel method for estimating the controllability Gramian matrix in a simple, theoretically grounded manner. This method provides insights into brain dynamics, including overall controllability (quantified by the Gramian’s eigenvalues) and specific controllable directions (represented by its eigenvectors). As a proof of concept, we applied our method to transcranial magnetic stimulation-induced electroencephalographic responses across four motor-related states and two resting states. We found that states such as open-eye rest, closed-eye rest, and motor-related states were more effectively differentiated using controllable directions than overall controllability. However, certain states, like motor execution and motor imagery, remained indistinguishable using these measures. These findings indicate that some brain states differ in their intrinsic control properties as dynamical systems, while others share similarities that make them less distinguishable. This study underscores the value of control theory-based analyses in quantitatively how intrinsic brain states shape the brain’s responses to stimulation, providing deeper insights into the dynamic properties of these states. This methodology holds promise for diverse applications, including characterizing individual response variability and identifying conditions for optimal stimulation efficacy.

**Significant statement:** The brain can be viewed as a control system transitioning between states, such as from rest to motor activity. Previous studies using network control theory mostly evaluated brain dynamics without external perturbations, neglecting their role in accurate system identification. This study integrates perturbation inputs with network control theory to propose a method for estimating the controllability Gramian, thereby providing insights into brain dynamics. We applied this approach to TMS-induced EEG responses in motor-related and resting states. Our findings show that controllable directions (eigenvectors) allow better discrimination between states than overall controllability. Our method can quantitatively assess brain state differences, and has potential applications in characterizing individual response variability and optimizing stimulation efficacy.

## 1 Introduction

Network control theory has been applied to neuronal networks to deepen our understanding of complex cognitive processes in neuroscience [Medaglia et al., 2017, Gu et al., 2015]. The approach models the brain as a control system, highlighting its ability to transition between states, such as from motor activity to rest. It introduces mathematically-grounded and interpretable measures to analyze brain activity as multidimensional time series across different regions. The core concept is controllability, which refers to the brain’s capacity to shift its dynamics from one state to another via internal or external inputs. Mathematically, controllability is defined using the controllability Gramian, a matrix that captures how the brain’s state changes under the influence of these inputs over time. This framework offers a more holistic understanding of neural processes compared with fragmented approaches focusing on specific activity features.

Pioneering studies have used network control theory to explore how the brain’s structural connectivity constrains its dynamics. They have provided mechanistic explanations for transitions between cognitive states, identifying key areas of the brain that can facilitate these transitions [Gu et al., 2015] and providing insight into optimal trajectories between states, such as moving from high activity in the default mode network to high activity in sensorimotor systems [Gu et al., 2017]. These studies have significantly advanced our understanding of how the structural connectivity of the brain governs its control capabilities.

Our study aims to advance the understanding of brain dynamics by shifting the focus from structural connectivity to dynamic neural activity data and examination of state-dependent changes in control properties. Previous research has largely relied on passive data (i.e., data without external inputs) [Li et al., 2023, Kawakita et al., 2022, Kamiya et al., 2023], limiting its ability to fully characterize the brain’s dynamic nature. To address this limitation, it is considered essential to apply perturbative inputs is essential for accurate system identification [Jakowluk, 2024, Ogino et al., 2025], enabling a more precise characterization of brain dynamics.

Although not explicitly grounded in control theory, the importance of perturbative inputs has been demonstrated in studies investigating cognitive states such as working memory [Rose et al., 2016], states of consciousness [Casali et al., 2010, Casali et al., 2013, Lee et al., 2022], and spatial attention [Okazaki et al., 2020]. However, while these studies underscore the value of perturbative approaches, their reliance on one-dimensional, global measures like the Perturbational Complexity Index (PCI) [Casali et al., 2010,Casali et al., 2013,Lee et al., 2022] limits the granularity of insights because these measures primarily capture the overall complexity or the degree of signal propagation in brain dynamics.

In this study, we integrate perturbation inputs with the control theory framework to introduce a novel method for quantifying multivariate state-dependent changes in network control properties. Our approach provides a simple yet theoretically-grounded method for estimating the controllability Gramian from neural activity data with perturbation inputs. The controllability Gramian offers insights not only into the overall controllability of brain dynamics, as quantified by its eigenvalues, but also into the specific directions in which these dynamics can be effectively influenced, as represented by its eigenvectors. Additionally, under the assumption that the perturbative input acts as an impulse, we show that standard control theory methods can be used to estimate the controllability Gramian directly from the time series responses alone, using standard control theory methods. This approach eliminates the need to estimate the system’s connectivity matrix *A* or input matrix *B* (see Eq. (1)), thereby simplifying the estimation process.

As a proof of concept, we applied our method to single-trial, single-pulse transcranial magnetic stimulation (TMS)-electroencephalographic (EEG) data, covering four motor-related task conditions and two resting-state conditions (Table 1). By treating the short TMS pulses (less than 2 ms) as impulse responses, our method allows the controllability Gramian to be directly estimated of the controllability Gramian from the TMS-evoked EEG data. This approach focuses on short-latency responses, minimizing the influence of peripheral effects, as demonstrated in previous studies [Conde et al., 2019, Biabani et al., 2021, Rocchi et al., 2021]. Even with such a short latency, the high temporal resolution of EEG allows for robust state-dependent characterization of neural dynamics. We demonstrate that our proposed method captures the state-dependent change in neural dynamics between motor-related and resting states in terms of both the magnitude and the directions of controllability.

**Table 1:** Experimental conditions.

| Condition | Abbreviation | Condition overview (TMS timing) |
| --- | --- | --- |
| Motor execution | ME | Right index finger isometric abduction for 3 s (1.5-2.5 s after the task onset) |
| Motor imagery | MI | Imagery of the motor execution task (1.5-2.5 s after the task onset) |
| After motor execution | AfterME | After the motor execution task (4-5 s after the task onset) |
| Resting state with open eyes | OE | (every 4-6 s) |
| Resting state with closed eyes | CE | (every 4-6 s) |
| Ipsilateral motor execution | iME | Left index finger isometric abduction for 3 s (1.5-2.5 s after the task onset) |

## 2 Materials and Methods

### 2.1 Theoretical background to control theory

In this study, we consider a linear auto-regressive (AR) model as a model of multidimensional brain dynamics and quantify the control properties of the system given some external input. Specifically, we consider the following model:

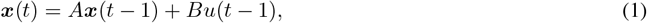

where ***x***(*t*) ∈ ℝ^*n*^ is a *n*-dimensional vector of the brain activity at time *t*,

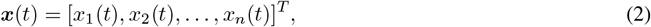

*A* is the *n*×*n* connectivity matrixthat describes the interaction between the elements, *u*(*t*−1) ∈ ℝ is a one-dimensional external input at time *t* − 1, and *B* is an *n* × 1 input matrix describing how the control input *u* affects the brain activity *x*. We consider the dynamics from the time *t* = 0 to *t* = *m* (Fig. 1).

**Figure 1.**
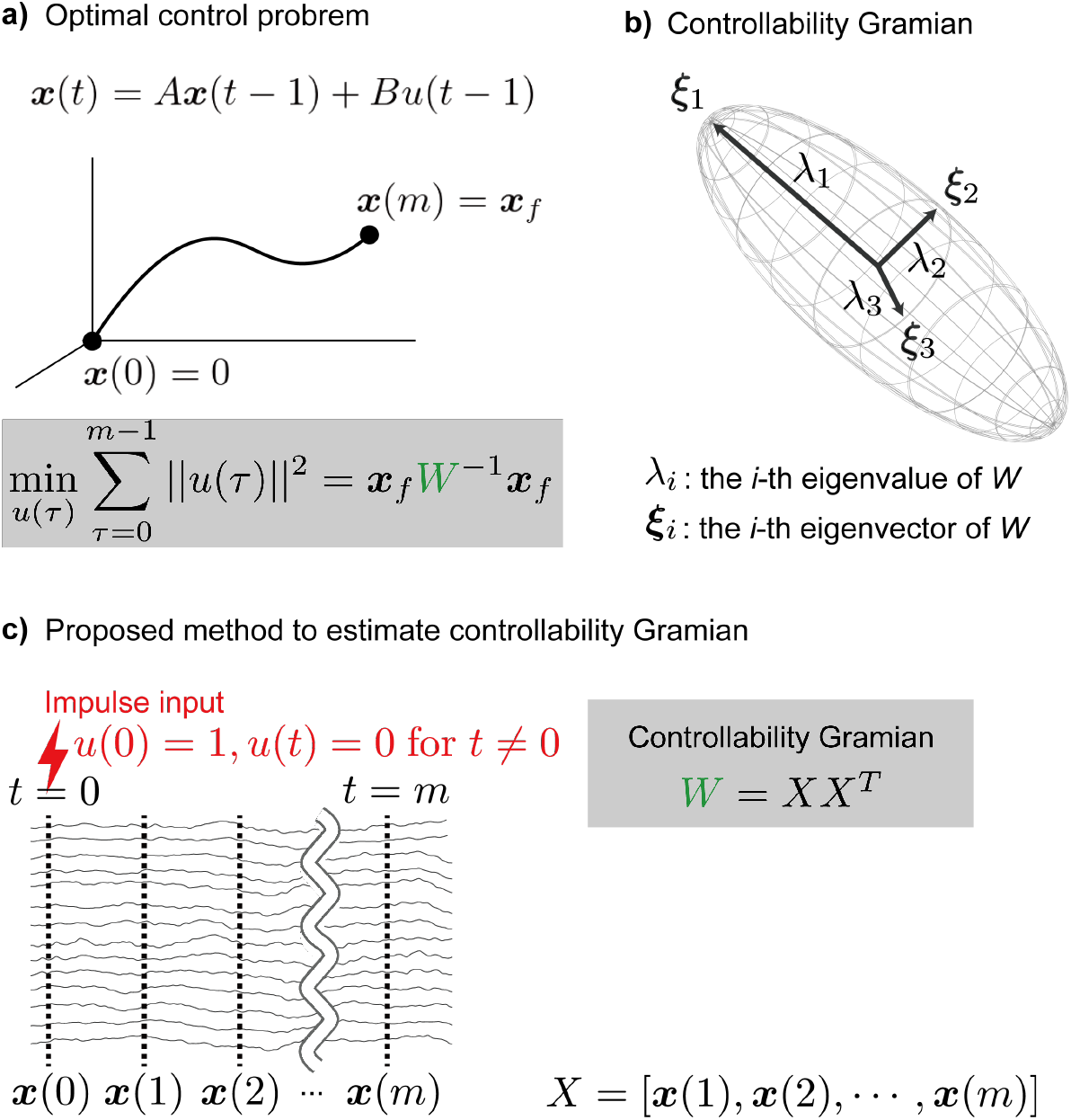
Schematic illustration of the controllability Gramian. (a) In a first-order auto-regressive model, the transition from the initial state ***x***(0) to the final target state ***x***(*m*) at minimum cost is realized if the inverse of the controllability Gramian is known, as the equation in the gray shaded box shows. (b) The Gramian *W* is an *n*-dimensional hyper-ellipse, encompassing all spatial patterns that appear in the controllability matrix. *n* is the measurement dimension as long as the measurement dimension does not exceed the number of time points. The major axis shows the first eigenvector ***ξ***_1_. The second and third eigenvectors ***ξ***_2_ and ***ξ***_3_ are shown as arrows orthogonal to the first eigenvector. The length of each axis corresponds to the magnitude of change as an eigenvalue *λ*., and the direction of a specific spatial pattern as an eigenvector ***ξ***.. (c) Assuming that the control input is an impulse, the controllability matrix can be regarded as the measurement signals *X*, as the equation in the gray shaded box shows. Under this assumption, we obtain the controllability Gramian without estimating the connectivity matrix *A* and input matrix *B*. Each line on the left indicates the time series measured by each sensor.

We wish to characterize both to what extent and how the external input *u* can control the system. To this end, the most important quantity to quantify is the so-called controllability Gramian. We will briefly explain the definition and meaning of the controllability Gramian, but for further details please refer to [Brunton and Kutz, 2019].

The controllability Gramian *W* is defined using the controllability matrix 𝒞 as follows:

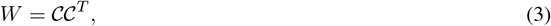

where

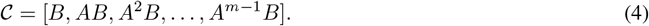

The controllability Gramian is important because it gives the solution to an example optimal control problem is: given the initial state ***x***(0) = ***x***_*s*_ and the final target state ***x***(*m*) = ***x***_*f*_, find the most efficient control inputs in terms of minimum cost.

Here, to simplify the expression, we introduce a new variable 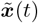, which is the time series ***x***(*t*) subtracted from ***x***(0), 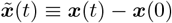, so that 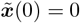 (Fig. 1a). Under this notation, the optimal control problem is formulated by

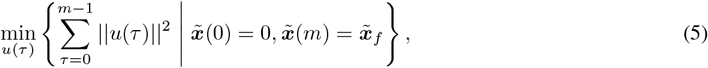

where 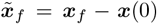. By solving this constrained optimization, we can find the minimal cost and the optimal control input according to:

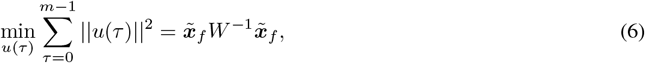

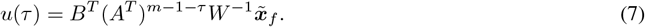

The equation for the minimum control cost (Eq. (6)) gives the exact mathematical meaning of the controllability Gramian *W* in terms of the optimal control framework. Assuming that the controllability Gramian has a set of eigenvalues *λ*_*i*_ and corresponding normalized eigenvectors ***ξ***_*i*_, then, from Eq. (6), we can easily see that the larger the eigenvalue is, the larger we can move the state of the system in the corresponding direction of the eigenvector. More precisely, given a fixed amount for the sum of the squared control input, i.e., 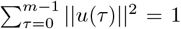, the squared distance we can move in the direction of an eigenvector is exactly given by its eigenvalue. This can be shown by setting the final target state to 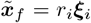, where *r*_*i*_ corresponds to the distance to move, and substituting this final state into Eq. (6) gives 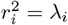.

In summary, by ordering the eigenvectors of the controllability Gramian according to the highest to lowest eigenvalues from highest to lowest, we can tell which direction we can easily move with the optimal control input: the direction of the largest eigenvalue corresponds to the most easily controllable direction, that of the second largest eigenvalue corresponds to the second most easily controllable direction, and so on (Fig. 1b). Note that the degree of ease of control is defined by the hypothetical optimal control input, not by the actual impulse input. In Fig. 1b, we illustrate this visually by considering the controllability Gramian as a hyperellipse, whose longest axis corresponds to the first eigenvector, whose second longest axis corresponds to the second eigenvector, and so on. In this study, we characterize the control property of the brain dynamics in this way.

### 2.2 Experimental design and statistical tests

#### 2.2.1 Participants

Seventeen healthy volunteers (11 women, age range: 18–29 years) participated in the experiment. All participants had slept between 6.5 and 11.0 hours the night before the experiment, were right-handed as assessed by the Edinburgh handedness inventory [Oldfield, 1971] (range: 65–100), did not smoke regularly except for one participant, had no history of neurological or psychiatric diseases, and had no contraindications to TMS. After receiving written and oral information about the experiment, the participants gave written informed consent before testing. All TMS-EEG data were collected between December 2016 and March 2017 as a part of a TMS-EEG project approved by the local ethics committee of the capital region of Denmark (H-15008824). The study was carried out under the guidelines of the Helsinki Declaration.

#### 2.2.2 EEG and EMG recording

EEG and electromyographic (EMG) signals were recorded with a NeuroOne Tesla system (Mega Electronics, Kuopio, Finland) with a 63-channel equidistant M10 EEG cap (EASYCAP, Herrsching, Germany) and surface EMG electrodes (Ambu Neuroline 710, Ballerup, Denmark). EMG electrodes were placed on the first dorsal interosseous muscle on the right hand, using a bipolar belly-tendon montage and a ground electrode mounted at the right ulnar styloid process. All EEG electrodes, including a ground electrode and a reference electrode of the cap, were prepared using NuPrep Skin Prep gel and ABRALYT HiCl EEG electrode gel (EASYCAP). EEG electrode impedance was maintained below 5 kΩ throughout the recording periods. EEG and EMG signals were digitized at 5 kHz with a 1250-Hz low-pass filter and DC filter.

#### 2.2.3 Neuro-navigated TMS-EEG

TMS was applied over the hand area of the primary motor cortex (M1-HAND) of the left hemisphere using a MagPro X100 with an MC-B70 figure-of-eight coil (MagVenture, Farum, Denmark). The coil was placed tangentially on the scalp, with the handle directed posteriorly at a 45^°^ angle to the sagittal midline. To reduce TMS artifacts in the EEG signals, a soft silicone rubber sheet (1.5-mm thick) was placed between the coil and the EEG electrodes. Biphasic pulses, in which the second phase induced a posterior-to-anterior current in the cortex, were employed. Continuous monitoring of the TMS coil placement was ensured via a frameless stereotaxic neuro-navigation system (Localite, Sankt Augustin, Germany). The root mean square of the disparity from the co-registered landmark over the M1-HAND was kept under 2 mm throughout the experiment.

The individual M1-HAND hotspot was identified as the coil position eliciting the most consistent and prominent motor-evoked potentials (MEPs) using suprathreshold stimulation intensity while the subject was at rest. This hotspot was marked in the neuro-navigation system. The resting motor threshold was determined after mounting the EEG cap using maximum-likelihood parameter estimation with the sequential testing approach while participants remained at rest [Awiszus, 2003]. MEPs exceeding 50 μV were classified as ‘responding.’

#### 2.2.4 TMS-EEG measurements

Participants underwent six conditions (Table 1), during which a single-pulse of TMS was applied to M1-HAND at a stimulation intensity of 120% of the resting motor threshold for each trial. One hundred trials were carried out per condition. Participants sat in a comfortable armchair, and auditory noise masking was consistently applied throughout the TMS-EEG measurements using in-ear headphones (Insert Earphone 3A 410–3002, 3M systems). Consistent with previous procedures, we employed specific time- and frequency-varying sounds for noise masking [Conde et al., 2019, Herring et al., 2015, Beck et al., 2024]. The sound pressure for noise masking was individually adjusted to a level where participants could not hear the click sound of the TMS pulse with the TMS coil placed on their M1-HAND or until they reached their upper threshold for comfort.

Measurements commenced with either the OE or CE conditions described in Table 1, with the order of these conditions being pseudo-randomized across participants. Participants placed their right hand, palm side down, on the armrest and remained at rest. In the OE condition, participants were instructed to keep their eyes open and fixate on a fixation cross, while the TMS pulse was administered every 4–6 s. In the CE condition, participants were instructed to keep their eyes closed, and the TMS pulse was delivered every 4–6 s.

Following the OE and CE conditions, three conditions involving right-hand movement or motor imagery (ME, MI, and AfterME) were conducted. In these three conditions, each trial began with the presentation of open gray and filled blue circles (ME and AfterME) or an open blue circle (MI) on a computer monitor placed in front of the participants. After the presentation of the two circles, participants performed 3 s of isometric abduction of the right index finger at 10% of maximum voluntary contraction. The size of the filled blue circle increased depending on the force level applied. Participants were instructed to match the size of the two circles as quickly as possible, meaning that the applied force level corresponded to 10% of the maximum voluntary contraction. In the MI condition, after presentation of the open blue circle, participants performed 3 s of kinesthetic motor imagery of the ME content without moving their right hand. After 3 s, a gray cross appeared on the monitor, which prompted participants to cease motor tasks and return to a resting state until the next trial began. TMS was administered 1.5–2.5 s from the start of each trial in the ME and MI conditions and 4–5 s from the start in the AfterME condition. The ME, MI and AfterME conditions were pseudo-randomized per trial and were paused multiple times to allow participants a short break and prevent mental fatigue.

Finally, we conducted the iME condition, in which participants were asked to perform the ME content with their left hand. TMS was administered 1.5-2.5 s from the start of each trial.

#### 2.2.5 EEG data preprocessing

Offline data analysis was performed using with the FieldTrip toolbox for EEG- and MEG analysis [Oostenveld et al., 2011]. Our proposed method compresses the time-series data in the temporal dimension (see Theoretical background to control theory). Since high-amplitude components can have lasting effects beyond their occurrence, the following preprocessing steps were implemented to minimize the impact of cranial muscle artifacts as much as possible. Initially, TMS-induced artifacts were removed from EEG signals using the TESA methods [Rogasch et al., 2017] with MATLAB software (MathWorks Inc., Natick, MA). In brief, the TMS-EEG data underwent the following processing steps: 1) removal of noisy channels; 2) epoching of data from −1 to +1 s relative to the TMS pulse for the OE and CE conditions and from −1 to +6 s relative to the onset of the motor task for the other conditions; 3) baseline correction with data from −500 to −110 ms relative to the TMS pulse for the OE and CE conditions or relative to the motor task onset for other conditions; 4) removal of periods containing TMS pulse artifacts and TMS-evoked muscle activity (−2 to 15 ms relative to the TMS pulse); 5) interpolation of missing data around the TMS pulse with constant amplitude values; 6) removal of noisy trials; 7) independent component analysis (ICA) and removal of components representing muscle artifacts; 8) replacement of data of the removed window (−2 to 15 ms relative to the TMS pulse) by interpolation using a cubic function; 9) second-order Butterworth filtering (1–80 Hz); 10) notch filtering; 11) replacement of data from the removed window with constant amplitude values; 12) independent component analysis to remove residual artifacts; 13) spatial interpolation of the rejected channels; 14) common average re-referencing; and 15) linear interpolation of the 0–20 ms signal was applied to completely eliminate the residual effects of artifacts. As a result of the above preprocessing, an average of 2.25 (SD 1.51, range 0 − 8) channels, an average of 13.99% (SD 12.10, range 0 − 58) of trials per session were excluded. Through these preprocessing steps, we obtained TMS-induced artifact-free EEG data.

#### 2.2.6 Extracting the controllability matrix from single-shot TMS-EEG

We then extracted the controllability matrix corresponding to Eq. (13) from the TMS-induced artifact-free data. Focusing on cortical responses, signals with short latencies up to 60 ms were included in the analysis, thereby avoiding peripheral responses, which have a strong influence on long latencies [Biabani et al., 2019, Rocchi et al., 2021]. The analysis was performed according to the following steps: 1) scaling by the standard deviation of signals from −200 to −5 ms before stimulation to correct for differences in signal values per electrode; 2) downsampling to 1070 Hz to align the number of time points with the number of channels; 3) baseline correction with the values immediately previous to satisfy the assumption that 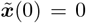; and 4) segmentation from 1 to 60 ms from TMS onset. This resulted in a 63-channel *×* 63-time-points data matrix of EEG signals for each trial. There is no need to estimate the connectivity matrix *A* and the input matrix *B* in Eq. (1).

When the analysis was performed over longer time windows, i.e., 1–120 ms and 1–300 ms, the downsampling frequency was set to 530 Hz and 220 Hz, respectively, so that the controllability matrix was a square matrix.

#### 2.2.7 Definition of pairwise distance

We treated the single-shot TMS-EEG data as the controllability matrix described in the previous section, from which we computed the controllability Gramians (Eq. (3)). For each trial, the resulting controllability Gramian was further analyzed through its eigenvalues and eigenvectors. To characterize trial-by-trial variations, we quantified pairwise distances between trials on the basis of these properties. This analysis was implemented in MATLAB using in-house code and the publicly available SPDtoolbox on GitHub (https://github.com/kyoustat/papers) [You and Park, 2021].

We used the following three distance measures.

##### Gramian-based distance

The controllability Gramians are considered to be on the symmetric positive definite (SPD) space, which is a Riemannian manifold. For any two SPD matrices *w* and *w*^′^, the affine-invariant Riemannian metric between them is defined as [Bhatia, 2009]:

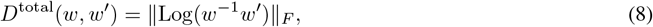

where Log(*a*) is the matrix logrithm of a square matrix *a*, and ∥*a*∥_*F*_ is the Frobenius norm of *a*.

When focusing only on up to the *m*th eigenmode (i.e., excluding the eigenmodes after the *m* + 1st eigenmode), the reconstructed controllability Gramian *ŵ* could be obtained accoding to the following. The original controllability Gramian *W* was decomposed into its eigenvalues and eigenvectors according to:

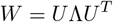

where *U* is the matrix of eigenvectors and Λ is the diagonal matrix containing the eigenvalues *λ*_1_, *λ*_2_, …, *λ*_*n*_ in descending order. To achieve dimensionality reduction, we reconstructed the Gramian matrix using only the top *m* eigenvalues *λ*_1_, *λ*_2_, …, *λ*_*m*_ and their corresponding eigenvectors. The reconstructed Gramian *Ŵ* is given by:

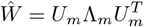

where *U*_*m*_ consists of the first *m* columns of *U*, and Λ_*m*_ is the diagonal matrix containing the first *m* eigenvalues. This allowed us to approximate the original Gramian in a lower-dimensional space.

##### Eigenvalue-based distance

Eigenvalues are considered as points on the complex number plane and Euclidean distance is used. The inter-trial distance of the average contorollability was calculated using the following equation:

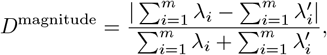

where *λ*_*i*_ is the *i*-th eigenvalue of one trial, and 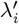 corresponds to that of another trial. *m* represents the number of eigenmodes, which is typically equal to the number of channels (*m* = *n* = 63) when all dimensions are utilized. However, *m* can be set to a smaller value than *n* when focusing only on the upper eigenmodes.

##### Eigenvector-based distance

The angle between the two eigenvectors was used to compare the direction independently of the magnitude. The distance between the *i*-th eigenvector of one trial *x*_*i*_ and the *j*-th eigenvector of another trial *y*_*j*_ was defined using the cosine distance as 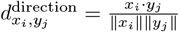. Using 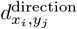, the distance of the controllable direction up to the *m*th eigenmode of a trial pair was determined as follows:

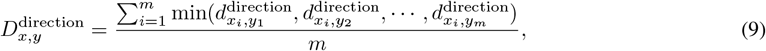

where *m* represents the number of eigenmodes. 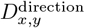 is the order-free cosine distance, which uses the distance of the most similar eigenvectors, regardless of the order of the magnitude of the eigenvalues. Since the resulting distance matrix is asymmetric 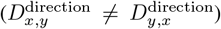, we symmetrized it by taking the arithmetic mean of each pair of corresponding trials, as follows:

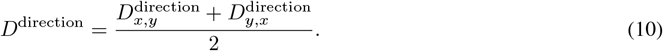

#### 2.2.8 Classification of conditions according to inter-trial distance matrices

Each of the three distance matrices we obtained used a different metric, and were thuson different scales and were difficult to compare. Therefore, we performed six-condition classification using the *k*-nearest neighbor method with each distance matrix and obtained a common index of classification performance (*k* = 5). Each trial was classified into one of six conditions, yielding a 6 *×* 6 confusion matrix. The confusion matrices were averaged across participants. Setting *k* to 1 or 3 and focusing on more local neighborhoods did not change the results.

#### 2.2.9 Measure for dimensionality of controllable directions

To verify the dimensionality of the controllability Gramian, the cumulative contribution rate up to the *m*-th eigenmode was determined as follows:

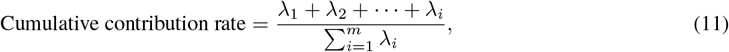

where *λ*_*i*_ is the *i*-th engenvalue. Rank, which is the number of eigenvectors with eigenvalues above a certain value (e.g., *λ*_⋅_ *>* 0.0001), is often used as another measure of dimensionality. When rank is used, factors such as measurement conditions (e.g., number of noisy channels) and preprocessing (e.g., how many independent components were excluded in the preprocessing) that are unrelated to the dimensionality of the controllable space and have no significant impact on the overall signal (eigenvalues are expected to be small) are affected. Therefore, in this study, dimensionality was evaluated using the cumulative contribution ratio, which allowed us to focus only on modes with large eigenvalues.

## 3 Results

Building on the theoretical background of control theory outlined in the Methods section, we first propose a method to estimate the controllability Gramian for evoked neural activity data acquired following an impulse response without explicitly estimating the system’s parameters *A* and *B*. The method itself is well known in standard control theory, but to our knowledge, it has not yet been used in neuroscience applications. Then, we demonstrate the utility of the proposed method by applying it to our TMS-EEG data. By integrating the proposed theoretical framework with empirical data analysis, we investigate the state dependence of the control properties of the EEG dynamics in response to single-shot TMS.

### 3.1 Proposed method to estimate the controllability Gramian

Here, under the assumption that the external stimulus is an impulse input, we propose a novel method to directly estimate the controllability Gramian, bypassing the need to estimate the system’s parameters. In general, to estimate the controllability Gramian from brain activity data with general external stimulation not restricted to an impulse input, we need to estimate *A* and *B* in Eq. (1). Once we have estimated *A* and *B* using some statistical methods, we can compute the controllability Gramian given by Eq. (3). However, in the special case where the external input is an impulse input, i.e., the input *u*(*t*) takes a non-zero value only at the time of stimulation (*t* = 0) and then takes zero values for the rest of the time ([*u*(0), *u*(1), … *u*(*m*)] = [1, 0, …, 0]), we do not need to explicitly estimate *A* and *B*, but compute the controllability Gramian directly.

The theoretical validity of this simplification can be demonstrated as follows. If we again introduce 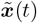 as in the Materials and Methods, and assume that the external stimulus is an impulse input, the time series of neural activity *X* is given by

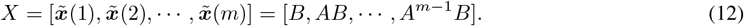

Comparing the above equation with the controllability matrix in Eq. (4), we can see that the time series of neural activity, *X*, is exactly equal to the controllability matrix 𝒞,

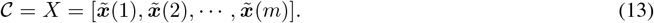

Finally, using Eq. (3), we obtain the controllability Gramian as

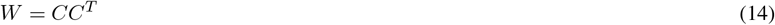

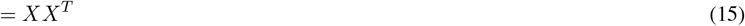

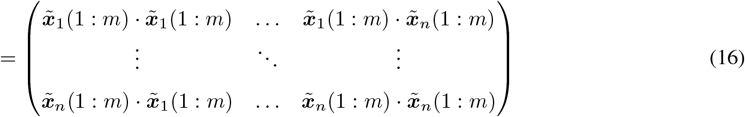

where 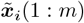 is the time series of the *i*-th element from *t* = 1 to *t* = *m*,

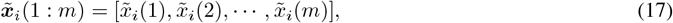

where 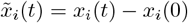 (Fig. 1c).

### 3.2 Examples of controllability Gramians from TMS-EEG responses

Using the proposed method illustrated in the previous section to compute the controllability Gramian, we here show several examples of controllability Gramian estimated from our TMS-EEG data. The analysis pipeline is summarized in Fig. 2a (see Methods for details). This study uses a dataset of EEG measurements made during TMS (Table 1). TEPs were recorded with a 63-channel EEG system (*n* = 63, see Materials and Methods). We set the time *t* = 0 to the time of TMS onset, and ***x***(*t*) is a 63-dimensional vector of TEPs at time *t* after the TMS onset. The TMS pulse is considered to be the impulse input, i.e., *u*(0) = 1 and *u*(*t*) ≠ 0 for any *t*≠0. An example of TMS-EEG response data acquired during the ME condition is shown in Fig. 2b. Then, using the equation for the controllability Gramian under an impulse input (Eq. (16)), we obtain an *n × n* controllability Gramian matrix from the EEG data after each single-pulse TMS. These matrices are illustrated in Fig. 2c, where the top and bottom panels show results from two different trials under the ME condition.

**Figure 2.**
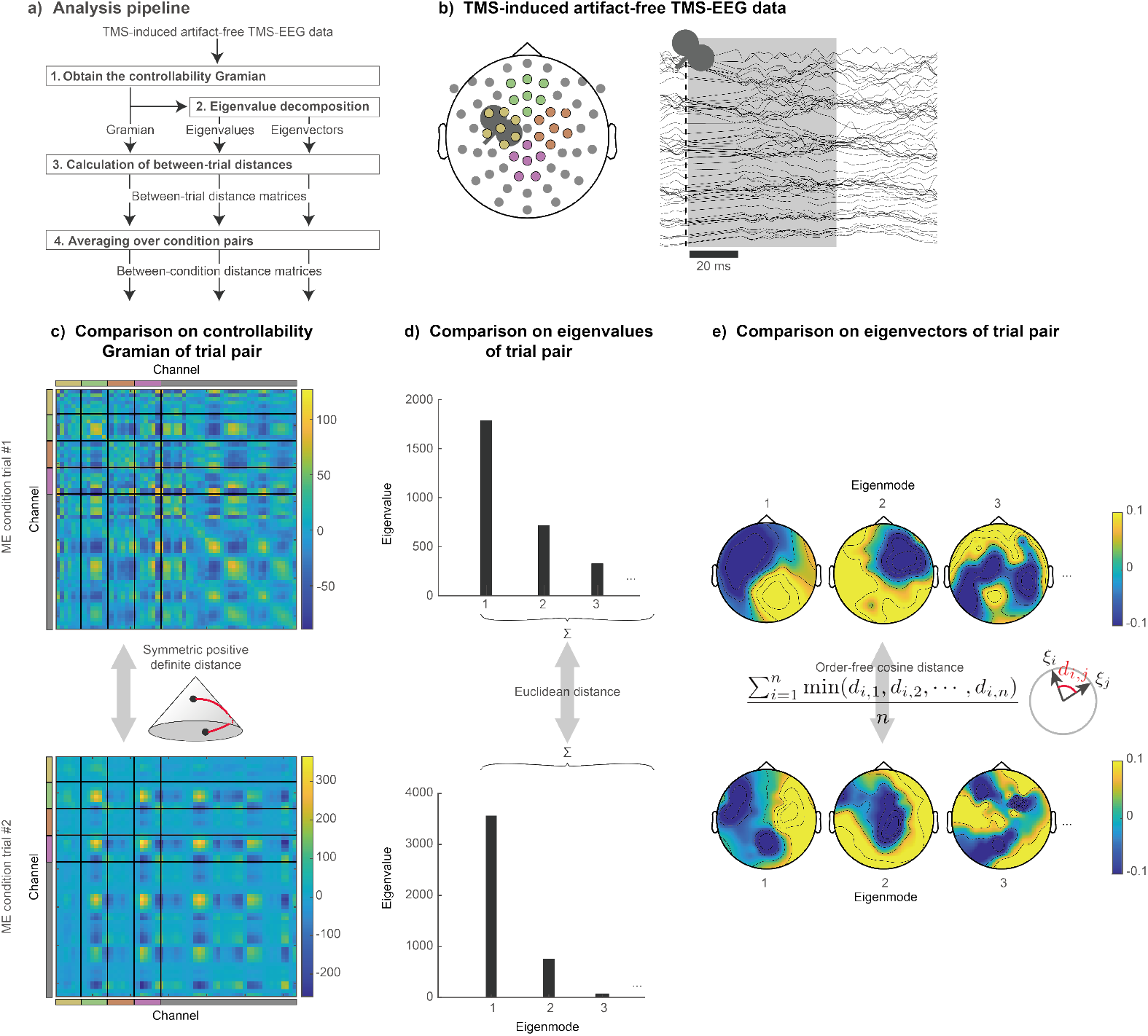
Overview of the proposed methodology and examples single trial analysis results. a) The controllability Gramian is derived from the single-trial TMS-EEG data after removing TMS-induced artifacts. Subsequently, the eigenvalues and eigenvectors of the controllability Gramian are computed. The comparative analysis entails the comparison of controllability Gramians and summation of their eigenvalues, and eigenvectors across each pair of trials to establish the distance matrices (see (c) and Methods). The average distance among trial pairs within the same condition is calculated to generate between-condition distance matrices. b) The left panel displays the electrode arrangement (with colors corresponding to (c)) and a sample waveform from a typical single trial. The temporal segments shaded in gray (1–60 ms from TMS onset) were used for the analysis. Note that the signal from 0 to 20 ms is linearly complemented to remove residual artifacts and the effects of artifact removal. c) The controllablity Gramians obtained from the EEG data of two different trials belonging to the same motor execution (ME) condition, are calculated for the distance in symmetric positive definite space. d) The Euclidean distance of the sums was used as the trial-to-trial distance for the eigenvalues. For the eigenvectors *ξ*, the effect of order defined by the magnitude of the eigenvalues was ignored, and only the directions were compared. e) Since eigenvectors are represented by relative values scaled between channels, each trial can be represented as *n* topographies. The distance between the *i*-th eigenvector of trial #1 *ξ*_*i*_ and the *j*-th eigenvector of trial #2 *ξ*_*j*_ was weighed by the cosine distance *d*_*i,j*_. The distance between trials was the average of the distances of the closest eigenvector pairs (see Methods). Note that the controllability Gramians, eigenvalues, and eigenvectors illustrated in c, d, and e were obtained by the proposed method applied to two typical trials.

### 3.3 Inter-trial distances based on the controllability Gramian

To explore the extent to which controllability characteristics vary among conditions, we computed the inter-trial distance based on the three metrics: the controllability Gramian-based, eigenvalue-based, and eigenvector-based distances (Figs. 2c–2e)(see Methods for details). As explained in the previous section, the controllability Gramian contains all the information for control properties associated with the TMS input. The distance in terms of the controllability Gramian represents the overall difference between trials in terms of control properties. By contrast, the distance for the sum of eigenvalues or eigenvectors represents the difference in terms of partial aspects of control properties, i.e., the overall degree of controllability and the direction in which the system can be controlled.

Figure 3a shows the inter-trial distance matrix based on the controllability Gramian. The indices of the matrix correspond to the trials of the TMS experiments, with each condition corresponding to one of six conditions: ME; MI; AfterME; OE; CE; or iME (Table 1). The trials are ordered from top to bottom according to the order of measurement for each session. The right-hand motor-related task conditions were performed in random order within the same sessions, while the other three conditions were performed in different sessions for each condition (see Materials and Methods for more details). Each session was arranged in the following order: the right-hand motor-related task conditions, i.e., ME, MI, and AfterME, and then OE, CE, and iME. The colors in the matrices represent the distance values, with blue colors representing low difference (i.e., the control properties of the trials are similar) and green to yellowish colors representing high difference, i.e., the control properties of the trials are different. We found that for all conditions, trials within the same sessions were close to each other, as indicated by the relatively small values of the elements of the diagonal blocks corresponding to respective sessions. For a more intuitive understanding of the distances between trials, we have converted the distance matrix shown in Fig. 3a to scatter plots in low-dimensional space with each trial as a point (Fig. 3b and Supp. Fig. S1). Dimension reduction using multidimensional scaling revealed that the two resting conditions (OE and CE) were clearly separated from the other motor-related conditions. In addition, iME was more localized, although it was mixed with the contralateral motor-related condition. It is worth noting that the clusters separated by these conditions span different sessions for each condition (Fig. 3c, numbers indicate sessions); that is, beyond intra-session similarity, clusters were identified by similarity between conditions.

**Figure 3.**
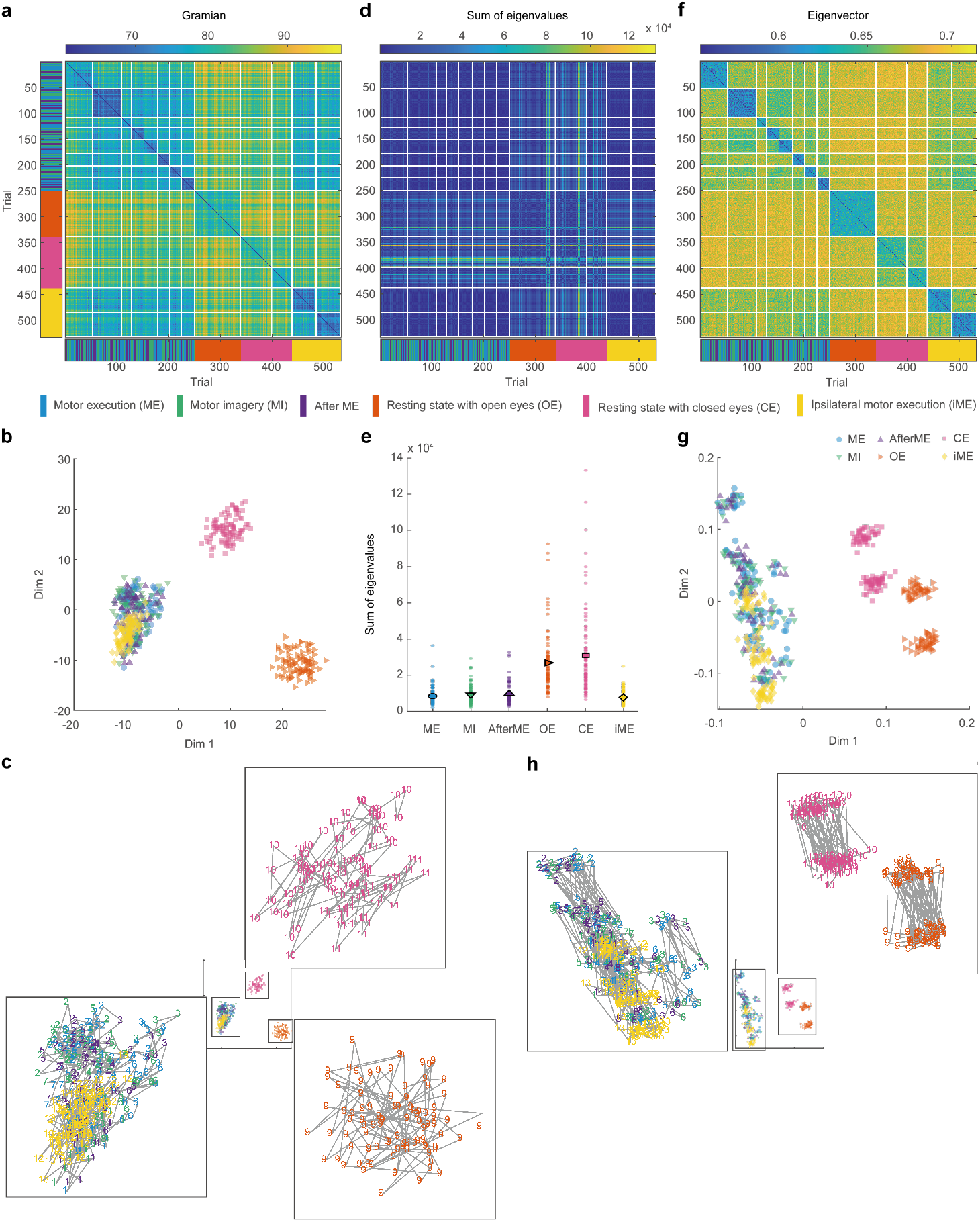
Distances between trials for controllability Gramians, sums of eigenvalues, and eigenvectors for a sample participant. a) A distance matrix of trials for Gramians. Each condition was measured in 100 trials, divided into 1 to 8 sessions. Note that the number of trials used in the analysis differs between conditions because of the noisy trial removal in the preprocessing. White lines indicate session boundaries and the color bar below indicates the condition to which each trial belongs in (b) and (c). b) The distance matrix illustrated in (a) reduced to a lower dimension using multidimensional scaling. Each point represents one trial, with colors corresponding to different conditions in (a). c) To clarify the distances within and between sessions in the same condition, the three clusters in (b) are enlarged. Numbers indicate session indices and colors indicate conditions. Consecutive trials are connected by gray lines. d) A distance matrix of trials for sums of eigenvalues. e) The sum of eigenvalues for each trial is represented as a dot, and the trial average is depicted as a colored symbol. f) A distance map of trials for eigenvalues. g) The distance matrix illustrated in (d) dimensionally reduced. h) The clusters in (g) are enlarged.

In a second step, with the sum of eigenvalues representing the controllable magnitude, we found that a distinct diagonal block structure in the distance matrix was absent, showing that trials within the same session were no closer to each other than trials from different sessions (Fig. 3d and Supp. Fig. S2), unlike situation for the controllability Gramian. While the sum of eigenvalues tended to be higher on average for the two resting conditions, there was considerable trial-to-trial variation (Fig. 3e). Thus, compared with the controllability Gramians, the sum of eigenvalues alone less well captures the differences between both conditions and sessions.

In a third step, using the eigenvectors we obtained a distance matrix more-or-less similar to that of the controllability Gramian (which is in contrast to the results for the sum of the eigenvalues), i.e., with the characteristic of small values in the diagonal block components of the distance matrix (Fig. 3f). In the reduced-dimensional space achieved by multidimensional scaling, the ability to distinguish between motor-related and rest conditions in the first dimension aligned with that of the controllability Gramians (Fig. 3g and Supp. Fig. S3). However, in the second dimension there was a wide spread of points representing motor-related conditions (cyan, green, purple, and yellow symbols), with no discernible difference from the resting-state conditions (orange and pink symbols). Even within the condition, the trials appeared to be split into multiple subclusters, similar to those found in the controllability Gramian. Both the average distance within a session (ME/MI/AfterME 0.59*±* SD 0.01, OE 0.61 *±* 0.02, CE 0.62 *±* 0.01, iME 0.59 *±* 0.01) and between successive trials (ME/MI/AfterME 0.60 *±* 0.01, OE 0.61 *±* 0.01, CE 0.63 *±* 0.01, iME 0.61 *±* 0.01) tended to be longer under the CE condition and a Friedman test revealed a significant effect of condition (within session *χ*^2^(37) = 3, *p <* 0.001, between successive trials *χ*^2^(28) = 3, *p <* 0.001). These suggest that one of the reasons for the wider distribution or more clusters of motor-related conditions than the resting state conditions is that data were collected in more sessions (ME/MI/AfterME 4.94*±*0.90, OE 1.28*±*0.39, CE 1.18*±*0.39, iME 1.94*±*0.56) rather than having large within-session or trial-to-trial variability. These subclusters were to some extent cohesive per session, but were not completely bounded. While they do not mix with each other in the OE, CE, and motor-related conditions, they do mix within the same condition across sessions, indicating a clear difference in controllable direction between conditions (Fig. 3h).

The following results were observed. First, there were differences between the resting and motor-related conditions in the Gramians, sum of eigenvalues, and eigenvectors. The sum of eigenvalues was larger in the resting state. Second, Gramians and eigenvectors formed multiple clusters in the lower dimensions. These clusters reflected differences between conditions, which were not solely due to slight variations in the measurements (e.g., electrode impedance and environmental noise), but also due to variations between the conditions. Third, the sum of eigenvectors showed that conditions were closer to each other across sessions. Subclusters were observed within the same condition, which were not observed in the Gramians. Fourth, the three motor-related conditions were more similar to each other than to the resting state conditions in terms of controllability, as indicated by their non-adjacency to the resting state condition trials in the low-dimensional space of Gramians and eigenvectors. While the influence of measurement states could not be completely eliminated, this finding suggests that the motor-related tasks shared common properties between the measurment sessions.

### 3.4 Inter-condition distances based on the controllability Gramian

To observe the average trend of the control properties across participants, we next quantified the averaged discriminability between conditions across participants using the distance matrices of the trials. We first averaged the inter-trial distance matrices for each participant across condition pairs and averaged the condition-based distance matrices across participants, obtaining Figs. 4a–4c. The distance matrices are difficult to compare with each other because different distance measures are used for the controllability Gramians, eigenvalues, and eigenvectors. Therefore, we performed six-condition classification using the *k* nearest neighbor method with each distance measure and obtained classification performance as a common measure (Figs. 4d, e, and f). Additionally, to better illustrate the differences and similarities between conditions, we constructed dendrograms from the distance matrices (Figs. 4g, h, and i).

**Figure 4.**
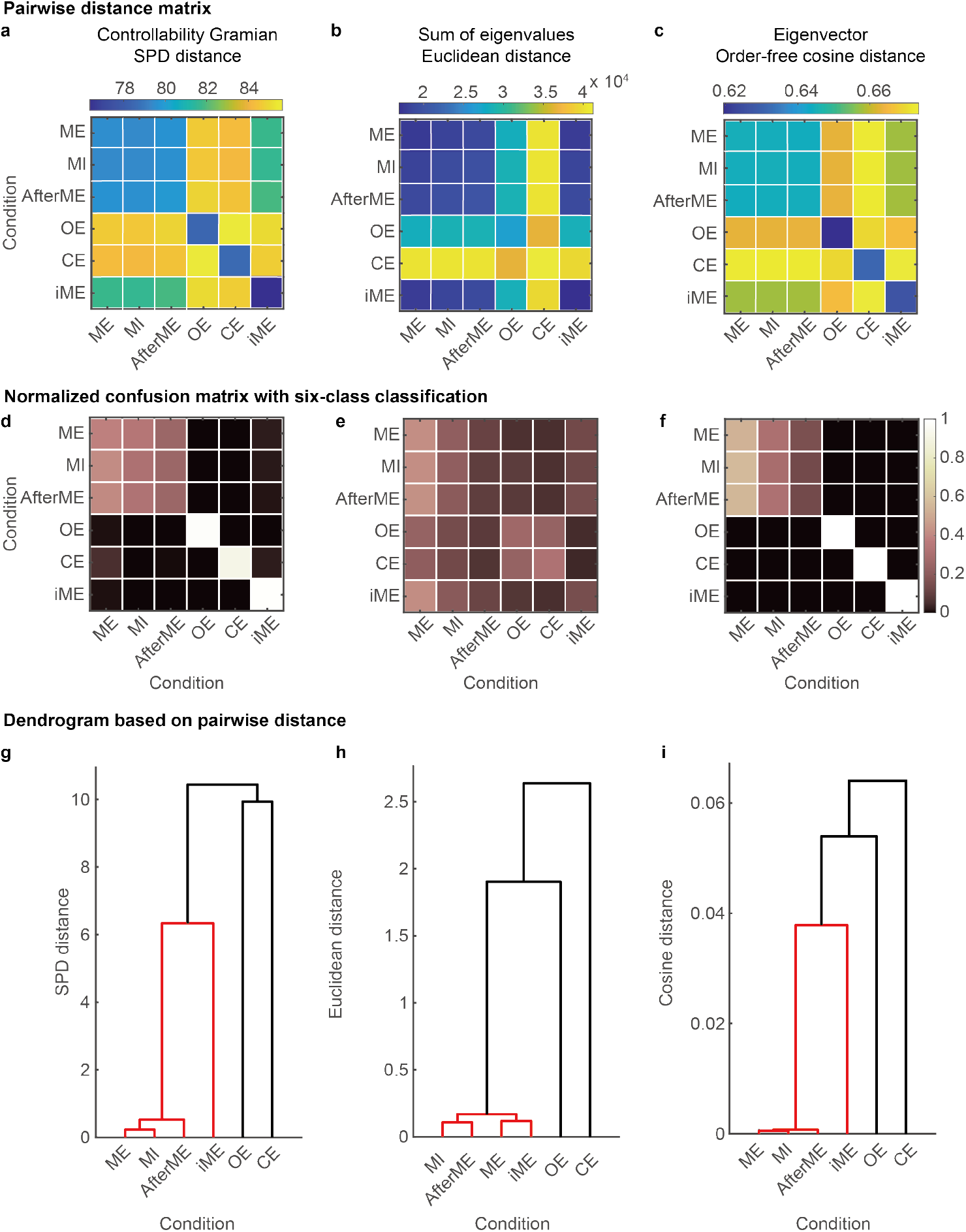
Distance between conditions for the controllability Gramians, sum of their eigenvalues and eigenvectors. a) Distance matrix between-condition pairs for controllability Gramians was averaged over 17 participants. b) Distance matrix between conditions for pairs sums of eigenvalues. c) Distance matrix between conditions for pairs of eigenvectors. d) Normalized confusion matrix for six-class classification using the *k*-nearest neighbor method (*k* = 5) in the distance space defined by (a). Rows represent the correct class and the columns the classified class. e) The normalized confusion matrix was obtained from (b). f) The normalized confusion matrix was obtained from (c). g) Dendrograms of the distance space defined by (a). Edges between node pairs that are less than 70% of the maximum linkage distance are indicated by red lines. h) The dendrogram obtained from (b). i) The dendrogram obtained from (c).

First, with the controllability Gramians, we found that the three contralateral motor-related conditions (i.e., ME, MI, and AfterME) were indisputably close to each other in the inter-condition distance matrices (Figs. 4a and d). The three conditions related to the right-hand motion were the most similar, with the left-hand motion condition being located between the resting conditions and the other motor-related conditions (Fig. 4g).

Second, using the eigenvalues, we found that the two resting conditions were distant from the other conditions, as was the case with the controllability Gramians (Fig. 4b). However, as seen in Fig. 3e, the variability between trials made it difficult to classify the conditions in each trial (Fig. 4e). The dendrogram based on the distance matrix between conditions showed that iME was most similar to ME, unlike the situation when using the controllability Gramians (Fig. 4h).

Third, using the eigenvectors, we found that in addition to the resting condition, which was clearly distinct from the other conditions in the controllability Gramians, iME also showed a trend toward a greater distance from the other conditions (Fig. 4c). In terms of classification performance, the three right-hand motor-related conditions were indistinguishable from each other, while the other conditions were not misclassified (as with the controllability Gramians) (Fig. 4f). The dendrogram showed that iME, like that of the controllability Gramians, was located between the three right-handed motor-related conditions and the two resting conditions, as was the situation for the controllability Gramians (Fig. 4i).

Taken together, the results suggest that controllable direction shows greater preservation of the discriminative ability of the controllability Gramians across conditions. Controllable magnitude represents a distinct aspect of variations in brain states to that represented by controllable direction. Ipsilateral and contralateral motion can be distinguished in terms of controllable direction, but not in terms of controllable magnitude. The three distances had one thing in common: four motor-related conditions including iME were closer to each other than 70% of the maximum distance (red lines in Fig. 4g, h, and i). Additional analyses were conducted and are presented in the supplementary material to address potential concerns regarding trial pairs within the same session and the impact of the time window for analysis. These analyses confirm that the observed results are robust, with the concerns identified concerns do not significantly influencing the outcomes.

### 3.5 Dimensionality of controllable directions

Finally, we examined the dimensionality of the space controllable by the TMS input using the cumulative contribution of the eigenvalues of the controllability Gramian. In short, when a small number of eigenmodes account for a high contribution rate, the dimensionality is considered low, indicating that a few dominant modes can effectively capture the data. By contrast, when a small number of eigenmodes show a low contribution rate, the dimensionality is high, suggesting that the data require a larger number of modes to be accurately represented. As illustrated in the controllability Gramians concept (Fig. 1b), high-dimensional controllability Gramians approximate hyperspheres, while low-dimensional ones exhibit a more flattened hyperellipsoidal form. For example, when the dimension is effectively 2 or 3, Gramians correspond to ellipses or ellipsoids.

We found that in all participants and in all conditions, 90% of the control properties elicited by TMS could be explained by the first to third eigenmode (Fig. 5a), indicating the low dimensionality of controllable space. Although there was no common trend among the participants as to which condition gave the highest contribution to the first eigenmode, we found that across a majority of participants, common trends emerged in contribution rates up to eigenmode 3: the contribution rate of CE was closest to 1, while that of OE was closer to 1 than the motor-related conditions, but there were no clear differences between the contribution rates of the motor-related task conditions (Fig. 5b). These results suggest that the controllability Gramian can be expressed in about 3 out of 63 dimensions in both resting and motor-related conditions, that resting states are lower dimensional than motor-related states, and that closing one’s eyes lowers dimensionality even more.

**Figure 5.**
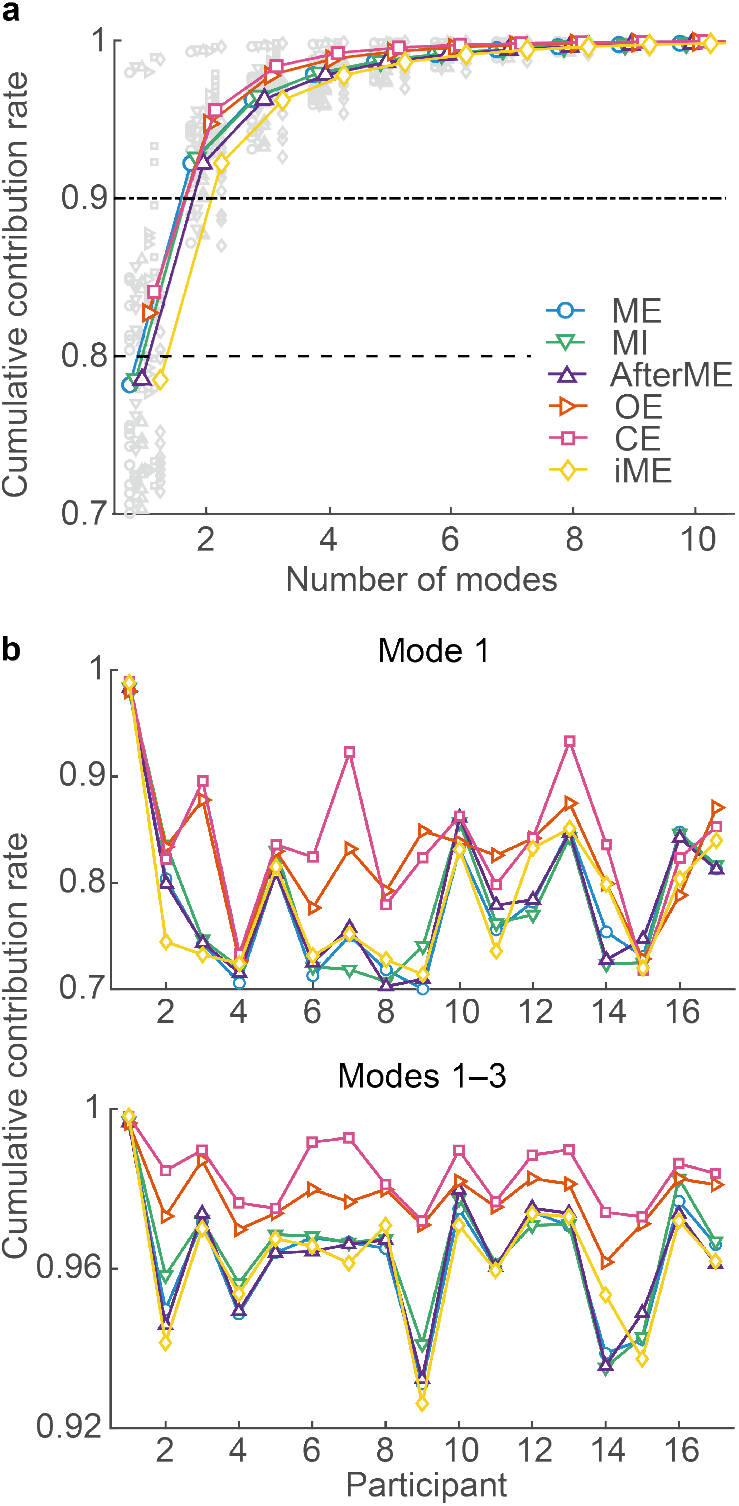
The control properties of M1 under TMS show high cumulative contribution rates for a small number of eigenmodes. a) Colored lines indicate across-participant averages of cumulative contribution rates. The values for each participant are indicated by gray symbols. b) Cumulative contribution rates for the first, and first to third modes. Color and symbols are the same as in a.

We additionally examined how these few large controllable directions relate to state dependence. Using only the top three eigenvectors, each trial was classified into six conditions, and we found that the correct rate was significantly lower than when all modes were used (Fig. 6). By increasing the number of eigenmodes used to 5, 10, and 30, the correct rate approached that when all modes were used. This suggests that even the lower eigenmodes, which do not provide a high contribution to amplitude restoration, retain information about the differences between conditions.

**Figure 6.**
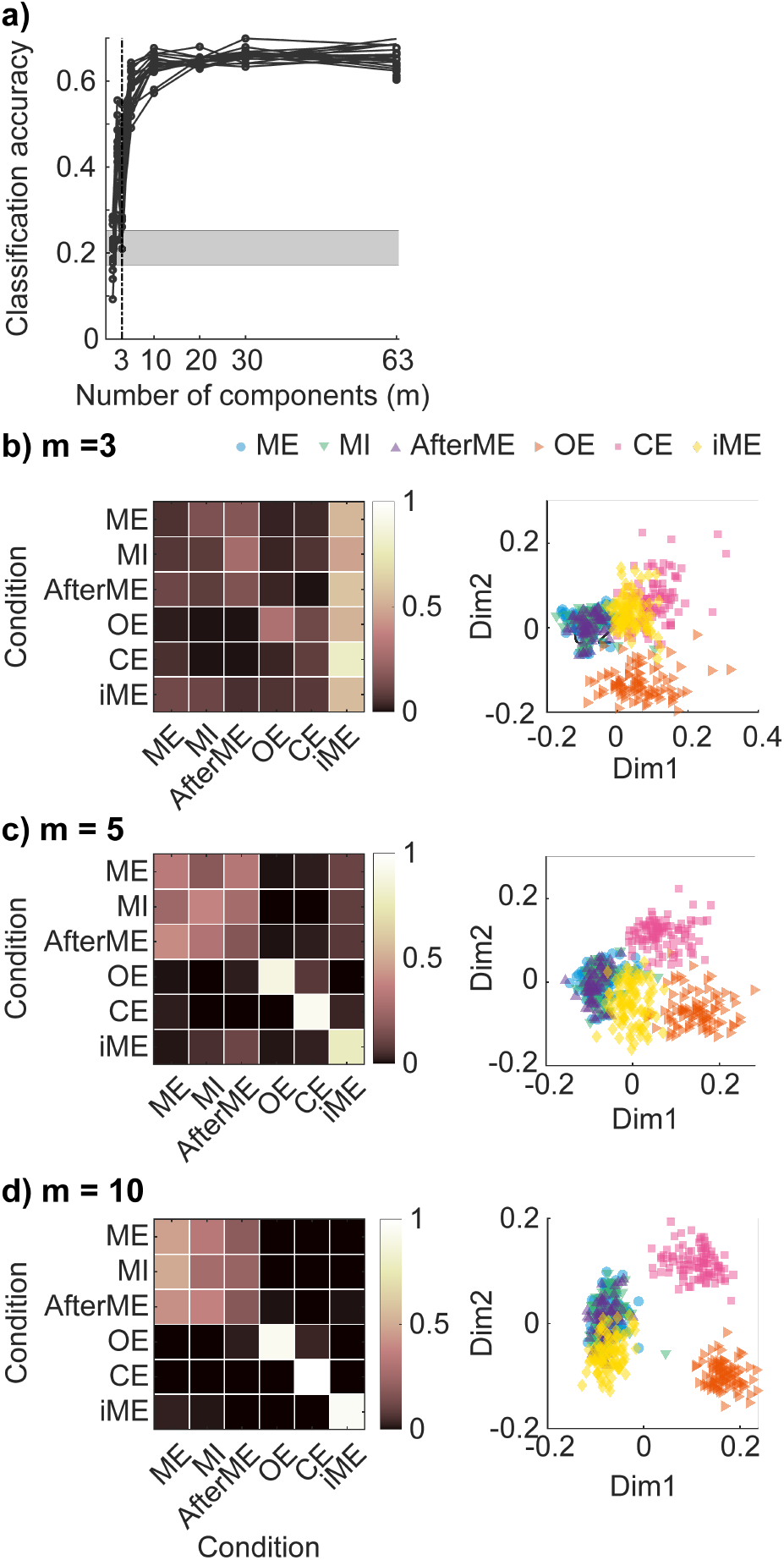
The performance of the six-value classification improved when a fourth or lower mode was added. a) The number of eigenmodes *m* and the percentage that were classified into the correct condition by the *k*-nearest neighbor method (*k* = 5). The black line represents the experimental participants, and gray shading represents the correct response rate obtained when predicting the most sampled condition (i.e., the chance level, the lower line represents the minimum between participants and the upper line represents the maximum). b) Confusion matrices (left) when using the first through to third eigenmodes, which account for 90% of the contribution. Classification performance was calculated for each participant and the participant average is shown. Distribution of trials in the classification space for one typical experimental participant (right). This three-dimensional classification space is represented in two-dimensions through the use of multidimensional scaling. Each point represents a trial. Confusion matrices and typical scatter plots when adding the mode used for condition classification from the first to the fifth (c) and first to the tenth (d) eigenmodes are shown.

## 4 Discussion

In this study, we propose a simple yet theoretically-grounded method based on control theory for analyzing the control properties of brain dynamics under impulse-like brain stimulation. On the basis of the standard results of the optimal control problem, we can regard the time series of neural activity as a controllability matrix, which enables us to simply compute the controllability Gramian without directly inferring the model parameters of linear dynamical systems. We applied our method to single-shot single-trial TMS-EEG data and used the controllability Gramian and its eigenvalues and eigenvectors to examine the differences and similarities in control properties across conditions. These control properties give a reasonable interpretation of responsiveness to local stimuli: the controllability Gramian represents the controllable space with unit input, its eigenvectors represent the controllable direction, and the corresponding eigenvalues represent the controllable magnitude. We found that controllability Gramians estimated from measured signals contains information that allowed distinguishing between open-eye rest, closed-eye rest, contralateral motor-related states, and ipsilateral motor execution state. This distinction is brought about by controllable direction rather than controllable magnitude. The cumulative contribution of eigenvalues indicated that the resting state had less response complexity, regardless of eye opening or eye closing. These results suggest that the state dependence of the EEG response immediately after TMS is characterized by spread rather than amplitude.

### 4.1 Novel perspectives of the proposed control theory-based measure on perturbation approaches

Although the proposed control theory-based measure is related to some measures quantified in previous studies, its main novel contribution lies in its ability to unify two aspects: the directions in which brain dynamics can be controlled and the extent to which they can be controlled with a fixed amount of input. In terms of relationships with previous studies, excitability and connectivity [Siebner et al., 2009, Bergmann et al., 2016] have previously been quantified as the magnitude of the response activity to a stimulus and the extent to which stimulus-evoked activity propagates [Casali et al., 2010, Casali et al., 2013, Casarotto et al., 2016, Comolatti et al., 2019]. The excitability and connectivity quantified in these previous studies are related to the sum of eigenvalues of the controllability Gramian in our framework, which quantifies the overall degree of propagation of the stimulus-evoked response. However, the eigenvector we focused on indicates the specific direction of this propagation. The novelty of our proposed method is that it quantifies the degree and direction of propagation represented by the eigenvalues and eigenvectors of the controllability Gramian, considering them as being completely separable. In previous explorations of control properties from measurement signals, only the eigenvalues were utilized and the corresponding eigenvectors were neglected [Gu et al., 2015], or they were used as eigenmodes without being separated [Comolatti et al., 2019]. In this study, we showed that differences s between conditions clearly appear in controllable directions rather than in controllable magnitudes.

### 4.2 Limitations of our controllability Gramian approach

The limitations of our network control theory-based approach stem from assuming a first-order linear AR model without noise. While this simplifies the controllability analysis and avoids the complexities of estimating connectivity (*A* in Eq. (1)) and input matrices (*B*), these assumptions are not fully realistic. First, in terms of model order, EEG responses often require higher-order AR models. Studies suggest an optimal order between 8.6 and 30, depending on the context [Tseng et al., 1995, Chang et al., 2012, Shakeel et al., 2021]. Extending the analysis to a higher-order model would more closely represent the real measurement data, but this would require estimating *A* and *B*, which are not needed in the proposed method. Second, regarding the linearity assumption, neural dynamics are in general nonlinear and thus there may be cases where nonlinear models are more appropriate. Note, however, that a previous study demonstrated that linear AR models remain reasonable approximations of macroscopic brain dynamics because of observational and technical limitations [Nozari et al., 2024]. Third, neural dynamics are in general stochastic, and thus accounting for stochasticity could change the prediction of control property models. In our previous work, we explored controllability in stochastic neural systems [Kamiya et al., 2023], and extending the current framework to incorporate stochastic systems is a potential avenue for future research, but this would substantially complicate the estimation of controllability. In addition, the proposed method is currently limited to impulse inputs; addressing more diverse input types would require explicit estimation of the connectivity and input matrices.

Despite these limitations, we found that the Gramian estimated using the simplified model effectively captures statedependent changes in controllability. A key advantage of this approach is that it simplifies the process by eliminating the need to estimate the connectivity matrix *A* and input matrix *B* (Eq. (1)), thereby enabling direct estimation of controllability Gramians. This simplicity makes the method especially suitable for researchers unfamiliar with the framework of control theory, providing a way to easily evaluate the control properties of complex brain dynamics that are typically difficult to assess. Future research should prioritize refining the estimation process by accounting for more realistic factors, such as nonlinearity and noise, to enhance the applicability and accuracy of our proposed method.

### 4.3 Validity of the results in comparison with previous studies

We found that the controllable magnitude fluctuates more during rest conditions than during motor-related conditions. One possible explanation is the reduced variability in brain activity induced by the visual stimuli used for task instructions, which stabilizes neural activity by constraining it to a specific trajectory [He and Zempel, 2013, He, 2011]. In addition, both motor execution and motor imagery enhance inter-trial coherence within the consistent activity of task-positive networks such as the somatomotor network [Churchland et al., 2010]. By contrast, during resting states, neural networks are less constrained by external demands, allowing greater flexibility and spontaneous variation between interconnected networks. This network-level flexibility may account for the higher trial-to-trial variability observed in resting-state EEG.

Our analysis further reveals that controllable directions retain more information for identifying states than do controllable magnitudes. Controllable direction is a relevant indicator of EEG spatial topography because it captures the relative relationships between EEG channels, rather than the absolute amplitude of each channel. Of note, the right-hand motor-related conditions and the two resting-state conditions formed distinct clusters in analysis based on controllable directions. Although the specific contributions of motor execution and imagery to these separations remain unclear, the distinction aligns with established findings, such as characteristic EEG topographies for visual evoked potentials and differences between open-eye and closed-eye resting states [Allison et al., 1977,Barry et al., 2007]. These findings confirm that motor-related states with visual instructions, open-eye resting states, and closed-eye resting states can be distinguished by their controllable directions.

We also found differences in motor execution between right and left hands. This is not surprising considering the stream of previous brain-computer interface studies taking advantage of topographic differences caused by differences in effectors [Xu et al., 2011, Huang et al., 2009], but it should be noted that our method does not take into account the sign of the eigenvector. In other words, the left and right-hand movements are not due to a simple lateral reversal of topography, suggesting that different neural activation patterns occur. The influence of afferent sensation on controllability could not be clearly concluded in this study; ME and iME could be distinguished, but not ME, MI, or AfterME. This may be related to the fact that motor imagery causes M1 excitability similar to that caused by motor execution [Takemi et al., 2013].

Taken together, our approach reveals that some brain states, such as motor-related task versus resting states, can be well distinguished by the properties of the controllability Gramian, whereas other states, such as motor execution and motor imagery, cannot be so differentiated. Future work is needed to verify which factors of brain states are reflected in the controllability associated with this TMS-EEG framework and which factors are not, with this involving performing TMS-EEG under more diverse conditions.

### 4.4 Considerations for Evaluating Control properties with TMS-EEG

In this study, the preprocessing steps used in TMS-EEG analysis could have influenced the results, particularly in terms of distinguishing between conditions. One such preprocessing method is ICA, which can affect the analysis by removing certain components. EEG signals consist of three types of components: (1) components that are common across states, (2) components that differ between states, and (3) noise. If components from category (2) are inadvertently removed, the ability to detect differences between states may be compromised. Therefore, it is crucial to carefully evaluate preprocessing methods to selectively extract components from category (2). Further research is needed to refine these approaches.

Filtering is widely used in EEG analysis and can significantly impact the results. While it helps reduce the influence of extracerebral sources, it may also eliminate signals that are critical for identifying specific states. Therefore, further refinement of preprocessing parameters is necessary to achieve a balance between selectively removing unwanted signals and preserving relevant state-specific information.

A key consideration for future research is the relationship between control properties and physiological indices. It is possible that the controllable direction is more closely associated with measures such as MEP, heart rate, and respiration rather than overall brain state, potentially leading to the emergence of subclusters within conditions. In this study, higher MEP values were observed in the ME, which is inherently linked to the other two motor-related conditions. This suggests that MEPs do not directly contribute to distinguishing between conditions but warrant further investigation.

Similarly, the relationship between controllability and traditional EEG features is also of interest. Given that TEP amplitude is a measure similar to controllable magnitude, it may have limited utility in differentiating between states. However, examining finer-grained information, such as latency, could provide deeper insights and should be explored in future studies.

## Acknowledgements

We would like to express our gratitude to Dr. Mikito Ogino for his valuable comments and suggestions on earlier drafts of this manuscript. We extend our heartfelt thanks to Dr. Ben D. Fulcher for his invaluable contributions and insightful discussions that greatly shaped the initial direction of this research. This work was supported by JST Moonshot R&D Grant Number JPMJMS2012 to M. O. and Y. S., JSPS KAKENHI Grant Numbers 18K15341 and 21K19810 to Y. S., the project “Precision Brain-Circuit Therapy - Precision-BCT” from Innovation Fund Denmark (grant nr. 9068-00025B) to H. R. S., the project “ADAptive and Precise Targeting of cortex-basal ganglia circuits in Parkinson’s Disease - ADAPT-PD” from The Lundbeck Fondation (collaborative project grant, grant nr. R336-2020-1035) to H. R. S., and Lundbeck Foundation Experiment grant (grant no. R346-2020-1822) to L. T.

## Supplementary Materials

### Inter-trial distance for all participants

In addition to the representative individual results shown in Figure 3, we here include a scatter plot of the Gramians (Supp. Fig. S1), their eigenvalues (Supp. Fig. S2), and their eigenvectors (Supp. Fig. S3) for all participants.

### Inter-condition distance excluding trial pairs in the same session

Consistency in non-task-related factors, such as impedance and environmental noise, was maintained within each session. This raised concerns that it might hinder the differentiation of the three contralateral motor conditions, whereas variability in the factors might facilitate such inter-condition differentiation. To investigate this possibility, distance matrices and dendrograms were generated after excluding trial pairs from the same session (Fig. S4). However, this manipulation did not alter the conclusions.

### Inter-condition distance including the TEP component for long latency

Another concern is the possibility that the short time window (1–60 ms) used in the analysis may have caused differences between conditions to be overlooked. To examine this possibility, distance matrices were generated with longer time windows (Fig. S5). The results showed that the patterns of the distance matrices were consistent, although the distances tended to get closer to inter-conditions. This is consistent with previous findings that show that the slow latency component of TEPs is associated with peripheral effects assumed to be consistent under conditions [Biabani et al., 2019, Rocchi et al., 2021].

**Figure S1:**
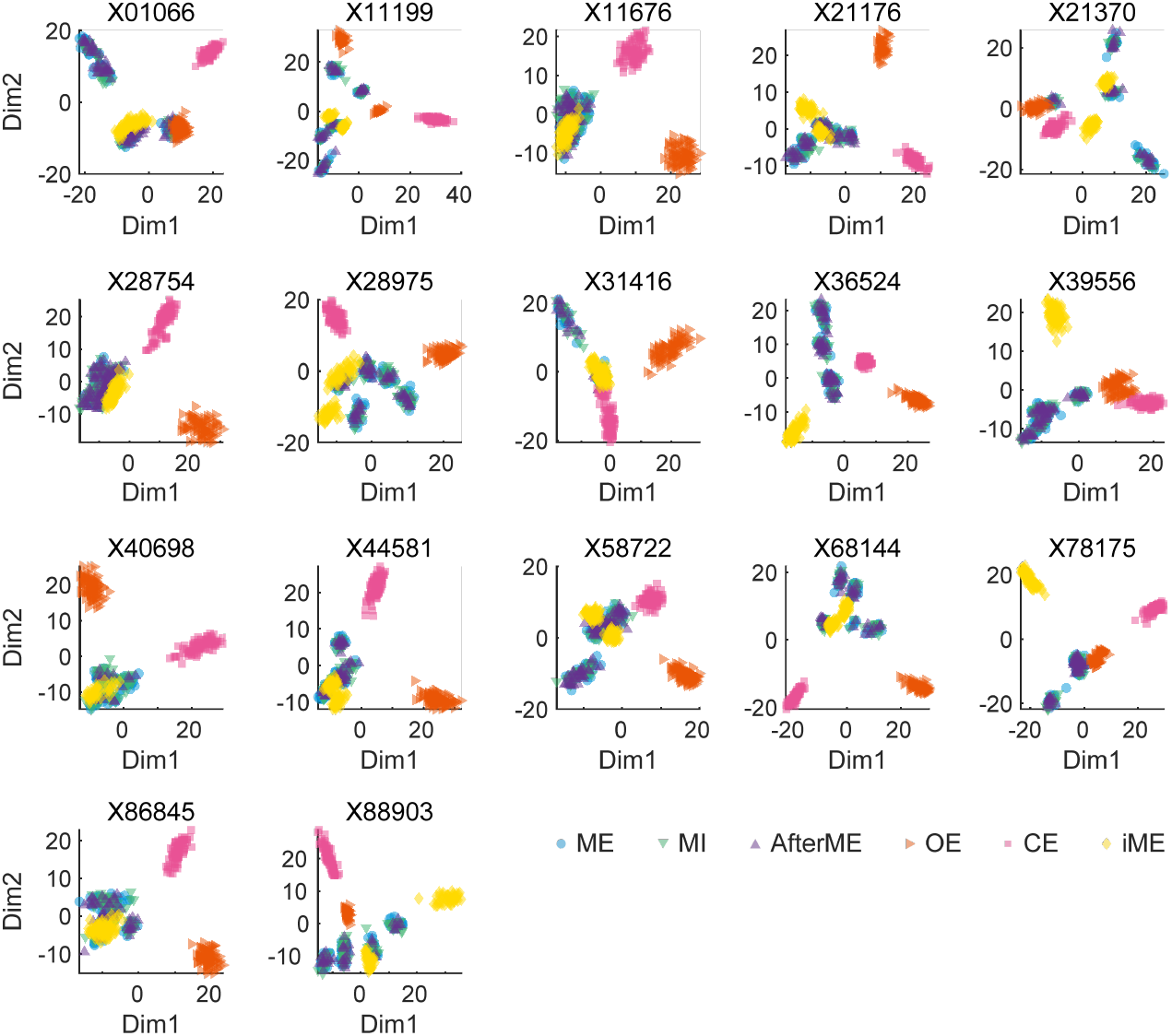
Two-dimensional representation of all trials from all participants using multidimensional scaling based on the controllability Gramians, as shown in Fig. 4b. The plot shows the relative distances between trials on the basis of their dissimilarity. The x-axis (Dim1) represents the first dimension and the y-axis (Dim2) represents the second dimension. Shorter distances between points indicate higher similarity, whereas larger distances indicate greater dissimilarity.

**Figure S2:**
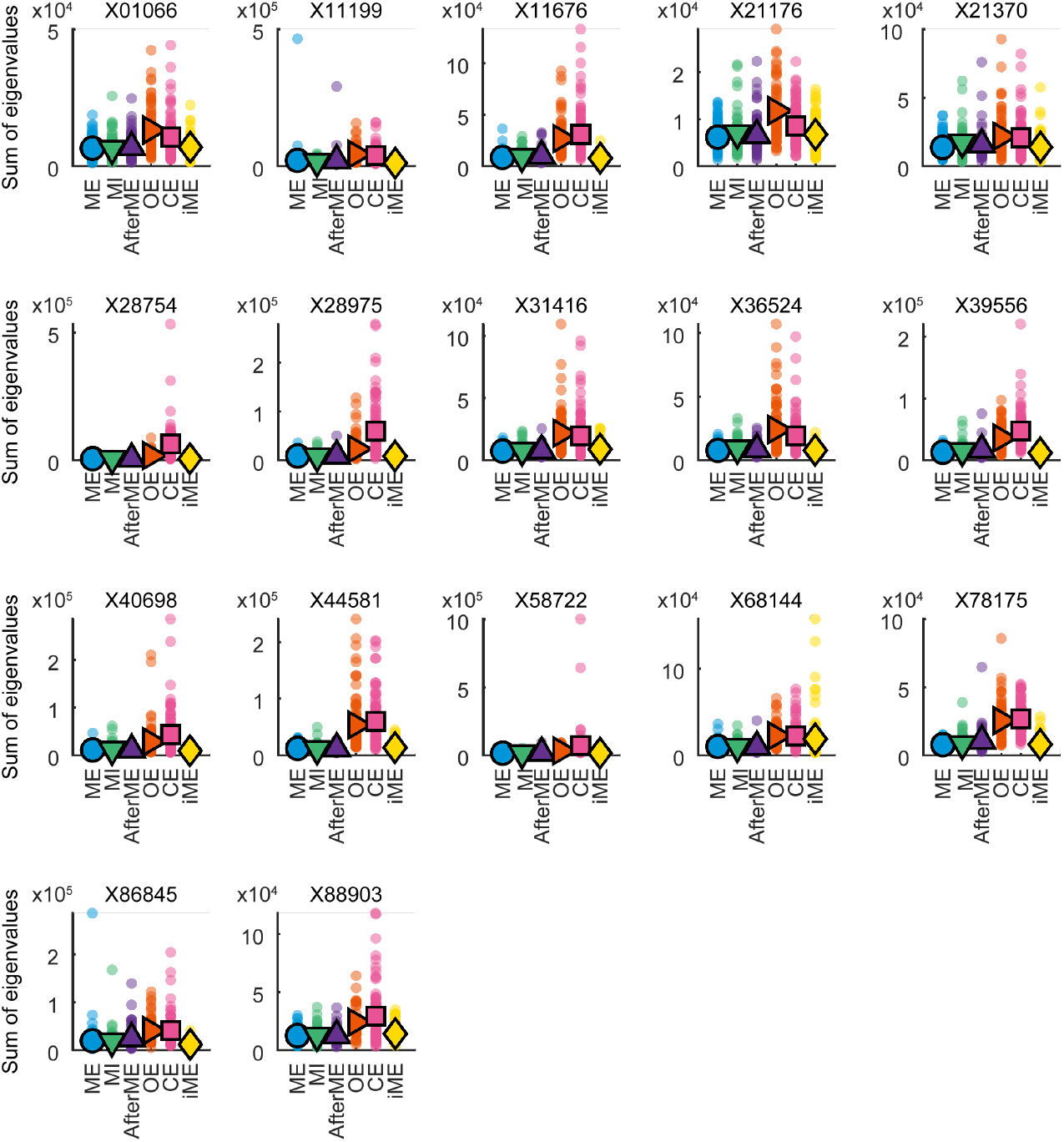
The sum of eigenvalues for each trial is represented as a dot, and the trial average is depicted as a colored symbol, as shown in Fig. 4e. Each panel indicates a participant.

**Figure S3:**
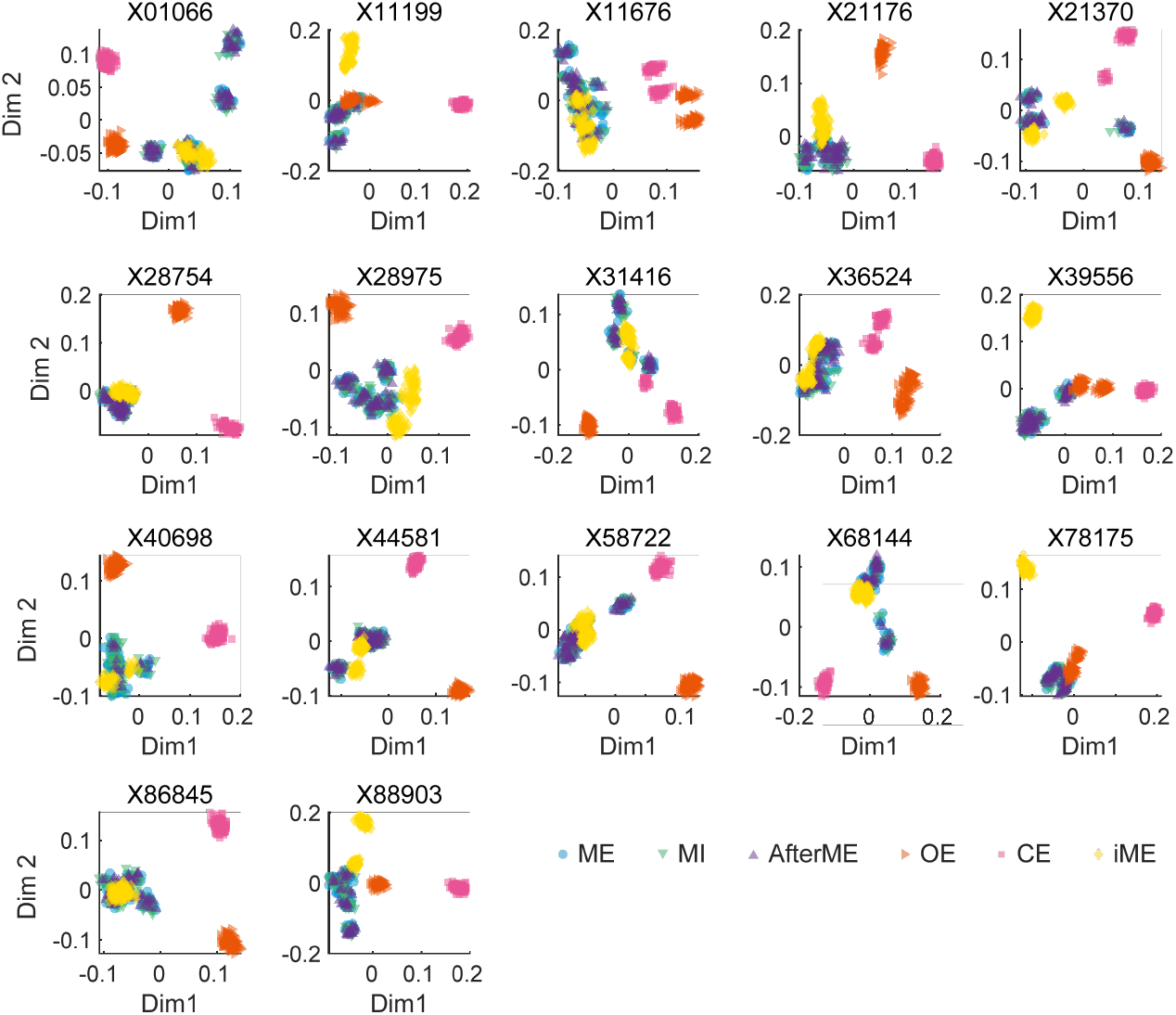
Two-dimensional representation of all trials from all participants using multidimensional scaling based on the eigenvectors, as shown in Fig. 4g. The plot shows the relative distances between trials on the basis of their dissimilarity. The x-axis (Dim1) represents the first dimension and the y-axis (Dim2) represents the second dimension. Shorter distances between points indicate higher similarity, while larger distances indicate greater dissimilarity.

**Figure S4:**
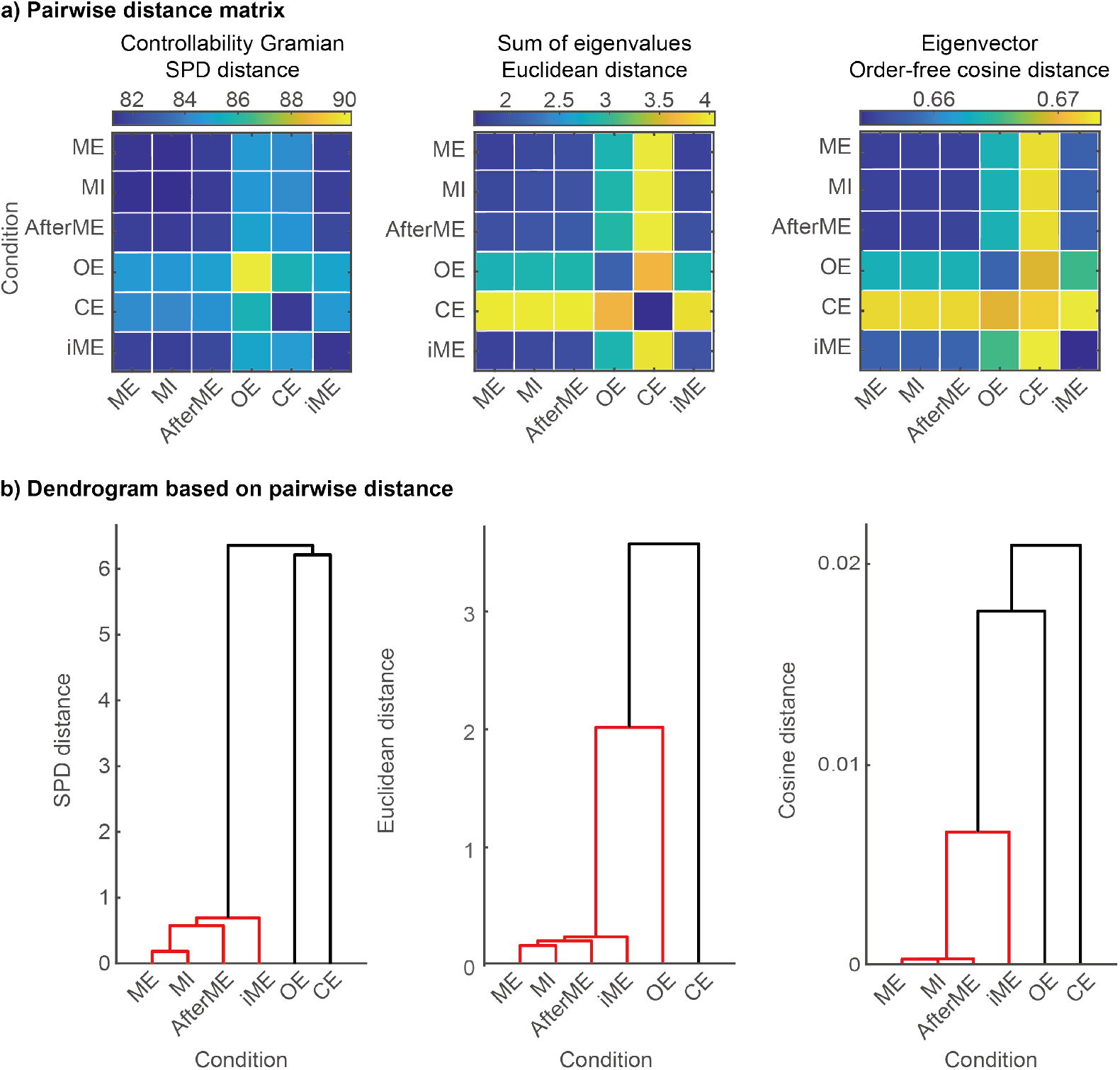
Distances between task conditions across sessions according to controllability Gramians, their eigenvalues, and eigenvectors. a) As in Figure 4, distances between trial pairs were calculated and averaged across condition pairs. However, samples belonging to the same session were excluded. Note that in the OE, CE, and iME conditions, the values of the diagonal components are unstable because multiple sessions were measured for only a few experimental participants. b) Dendrograms in the distance space defined by (a). Edges between node pairs that are less than 70% of the maximum linkage distance are indicated by red lines.

**Figure S5:**
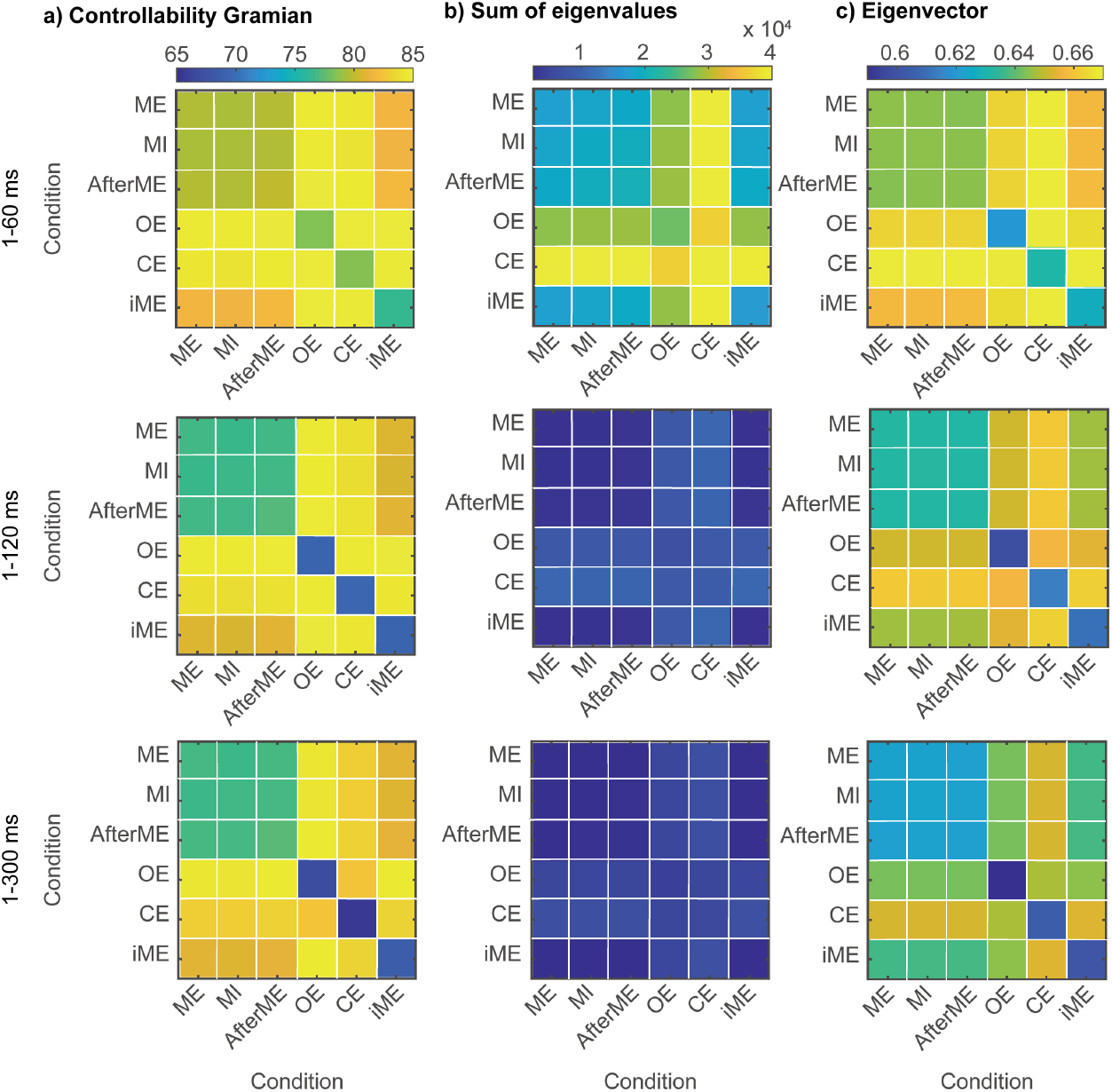
Longer time windows reduce distances between conditions. Distances between conditions when time windows are 1-60, 1-120, and 1-300 ms. The upper row is identical to Figure 4a, but the color scale has been adjusted for clarity. a) Distance matrices of controllability Gramians. b) Distance matrices of sums of eigenvalues. c) Distance matrices of eigenvectors.

